# Fast retrieval of structurally similar antibodies from large sequence databases with AbSLang

**DOI:** 10.64898/2026.08.05.742817

**Authors:** Eric Ji Da Wang, Fabian C. Spoendlin, Alexander Greenshields-Watson, Christopher R. Taylor, Charlotte M. Deane

## Abstract

The first steps in antibody therapeutic discovery involve identification of sequences with desirable binding properties. A way of finding these lead molecules is through the search of large sequence databases. Current methods, due to the size of databases, rely on germline or complementarity-determining-region (CDR) sequence identities, overlooking structurally similar antibodies with divergent sequences which can have identical binding properties . To address this, we introduce AbSLang, a model trained for pairwise CDR RMSD prediction using a contrastive learning approach. We demonstrate that AbSLang has comparable accuracy to exact RMSD calculation after explicit structure prediction with state-of-the-art models. Building on this model, we implemented AbSLang-search, a pipeline for retrieval of structurally similar antibodies from large sequence databases. AbSLang-search is highly compute efficient and allows to search datasets with 10 million sequences in less than 2 seconds.

## 1 Introduction

Antibodies serve as the primary protective mechanism in viral vaccines [1] and are a rapidly expanding class of biologic therapeutics [2].

Antibody specificity is primarily driven by six hypervariable complementarity determining region (CDR) loops (three on heavy and three on light chains) on two variable domains (VH and VL) that directly engage antigens [3, 4]. Through V(D)J recombination and somatic hypermutation, the human immune system generates billions of unique antibodies capable of recognizing a vast array of epitopes [5, 6, 7].

Advances in high-throughput single-cell sequencing now generate massive datasets of paired VH-VL sequences, providing deep insights into immune dynamics [8, 9, 10]. Currently, the predominant computational strategy for categorizing antibodies into functional groups is ‘clonotyping,’ which clusters antibodies originating from the same progenitor B-cell lineage [11]. Clonotyping typically requires identical heavy chain V and J genes, matching CDRH3 lengths and *≥* 80% CDRH3 sequence identity. Clonotyping is mostly carried out on the heavy chain only, as requiring matching heavy and light chains can be too restrictive [12] and excludes many antibodies which are functionally similar. While clonotyping is highly accurate at identifying antibodies with shared binding properties, it fails to detect the full breath of functional equivalence as antibodies originating from distinct genetic backgrounds can adopt highly similar structures with similar physicochemical properties and recognize the same epitope [13, 14, 15, 16]. This structural convergence reflects functional similarities that sequence-based methods alone often fail to capture, highlighting the necessity of incorporating structural analysis into antibody characterization.

For example the SPACE2 method [15, 16] clusters antibodies based on the CDR RMSDs of predicted models, allowing it to detect antibody functional convergence beyond clonal lineage. However, such approaches based on explicit protein structure prediction remain computationally intensive, requiring significant resources to predict and store structures and to perform pairwise RMSD calculations, limiting their practical application to large-scale antibody sequence databases.

Structural similarity and homology of proteins can be predicted more rapidly from their sequence using the embeddings of specialized protein language models (PLMs). For example TM-vec [17] accurately predicts TM-scores [18] between sequence dissimilar proteins [17]. However, TM-vec is not suited for antibodies. Firstly, the method has not been trained on large amounts of antibody data. Secondly, TM-scores are a poor metric to compare the structural similarity of antibodies as the conserved immunoglobulin domain results in consistent high scores.

Here, we introduce AbSLang, a pair of models (one for VH and one for VH/VL) predicting CDR RMSD. The models were trained using contrastive learning to produce sequence-level embeddings where the vector similarity of any two antibodies reflects their structural similarity. We also release equivalent models for camelid heavy chain only antibodies (see SI). We demonstrate that AbSLang has accuracy comparable to predicting structures with state-of-the-art protein structure predictors followed by exact RMSD calculation while maintaining more than 100 times faster speed and more than 500 times lower memory usage. Building upon AbSLang, we implement AbSLang-search, a pipeline to retrieve structurally similar antibodies from sequence databases magnitudes faster than previous available methods making it feasible to search databases as large as OAS (2.5 million paired and 2.8 billion unpaired sequences) [19]. AbSLang and AbSLang-search are available on Github (github.com/oxpig/AbSLang) and through a web app which allows comparison of pairs of antibodies as well as searching OAS by structural similarity (www.opig.stats.ox.ac.uk/webapps/abslang).

## 2 Materials and methods

### 2.1 Model

We trained two models to predict the structural similarity of antibodies, one for VH and one for the full Fv (VH/VL pairs). An additional model for VHHs is presented in the SI. Our approach, inspired by TM-vec [17], takes residue level PLM embeddings as input and produces a 512-dimensional sequence-level representation.

AbSLang uses a twin neural network architecture. The backbone consists of 8 transformer encoder layers with FlashAttention [20], followed by a weighted-pooling operation that aggregates residue-level representations into a single fixed-length vector. This pooled vector is then projected by an MLP head into a 512-dimensional embedding. The network is trained so that the Euclidean distance between a pair of embeddings approximates the average CDR RMSD of the corresponding antibody pair; concretely, we minimize the L1 loss between the predicted (Euclidean) and target (RMSD) distances.

Structural similarity is quantified by calculating the average root mean squared deviation (RMSD) of the antibody’s CDRs. After aligning antibody structures based on their framework region *C*_*α*_ coordinates, the RMSDs between CDRs are computed using dynamic time warping (DTW) [21] to allow comparison between CDR loops of different lengths.

Input PLMs were chosen based on validation performance. The paired VH/VL model uses IgT5 [22]. The heavy-chain only and VHH models use ESM-C [23].

All training was performed using the AdamW optimizer with a CosineAnnealingLR scheduler. Each model is trained for a maximum of 50 epochs on 2 RTX6000 Ada GPUs with a learning rate of 1e-4 and a batch size of 256. Weights were selected based on lowest validation loss. When benchmarking computation time, all tests were run on a single RTX6000 Ada GPU, for index speed comparisons a single Intel(R) Xeon(R) Gold 5415+ CPU was used.

### 2.2 Data

Models were trained using structures from the Structural Antibody Database (SAbDab) [24]. The training, validation and test splits replicate those used by ABodyBuilder3 [25], ensuring that no sequences in the validation or test sets share identical CDR sequences with any sequences in the training set. The dataset contains 8,645 antibody structures (8375 training structures, 150 validation, 100 test). As the model was trained and evaluated as a twin model, train/validation/test sets consist of all possible pairwise combinations of structures within their respective data splits. (See SI for full lists of test set PDB IDs)

For comparisons against AlphaFold3, we used a subset of 33 antibody structures from the ABodyBuilder3 test set. These structures were released after the AlphaFold3 training cutoff date (September 30th 2021) and were filtered to remove CDR sequences identical to those in AlphaFold3’s training set (see SI for the list of PDBs).

### 2.3 Antibody sequence database search

We leveraged FAISS [26] to search for embedding similarity within large antibody sequence databases. We store model output embeddings in a FAISS index by L2 (euclidean) distance, allowing retrieval of top-K database sequences with the lowest CDR RMSDs. We implemented indexing in IndexPQ (product quantization) for large databases such as paired-OAS [19] (2.5 million unique VH/VL pairs) to reduce storage requirements.

### 2.4 Model evaluation

Model performance was benchmarked against ABodyBuilder2 [27], ABodyBuilder3 [25] and AlphaFold3 [28]. ABodyBuilder2 and ABodyBuilder3 were run according to the authors’ instructions. AlphaFold3 was run via the webserver (alphafoldserver.com). For heavy-chain-only benchmarking, paired sequences were modeled using ABodyBuilder2, ABodyBuilder3, and AlphaFold3, after which the resulting light chains were removed to focus exclusively on heavy-chain structures.

Raw RMSD prediction between antibody pairs is evaluated based on correlation to the ground-truth pairwise RMSDs using Pearson’s R and absolute error.

We also evaluated the model’s accuracy in identification of the most structurally similar antibodies, using each sequence in the test set as a query sequence and searching against the rest of the test set, specific metrics include:

- Precision in identifying structurally similar pairs: accuracy in identifying all antibody pairs under 2 Å in average CDR RMSD.
- Top-K retrieval: We evaluated the model based on retrieving the top 20 most structurally similar antibodies for each query. Overlap: number of antibodies appearing in both the predicted top 20 and the true top 20, averaged across all query antibodies. Rank correlation (Kendall’s *τ*): the similarity in the ordering between the predicted and ground-truth lists.

## 3 Results

### 3.1 Accurate prediction of CDR RMSD

We develop AbSLang for the prediction of Fv (VH/VL) and VH CDR RMSD across pairs of antibody sequences. The models produce a structure-aware vector embedding for each antibody which was trained that the Euclidean distance between two vectors represents the CDR RMSD of the corresponding antibodies.

We evaluate against an alternative approach using explicit structure prediction with antibody specific (ABodyBuilder3) and general protein (AlphaFold3) models followed by exact RMSD calculation. Both AbSLang models show comparable CDR RMSD prediction performance to ABodyBuilder3 and AlphaFold3-based workflows across the test set (Figure 1) measured by the correlation between true and predicted RMSD and mean absolute error (MAE) in the predictions.

**Figure 1.**
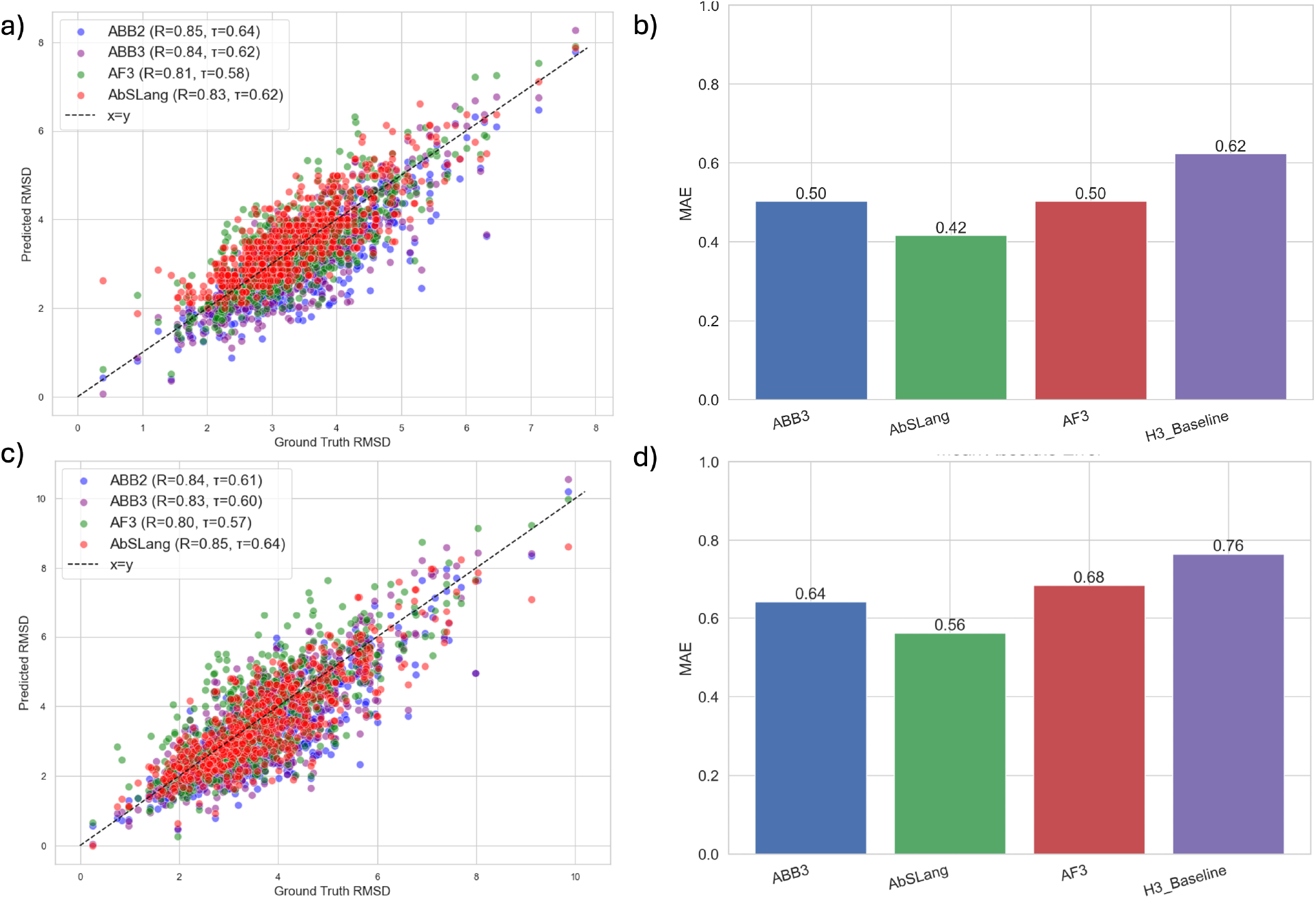
CDR RMSD prediction accuracy of paired VH/VL AbSLang (a, b) and heavy-chain only AbSLang (c, d). The scatter plots in the left column (a, c) show the ground truth pairwise CDR RMSD on the x-axis and the predicted RMSDs on the y-axis. Correlation between ground truth and predicted CDR RMSDs for AbSLang, ABodyBuilder2/3 (ABB2/3) and AlphaFold3 (AF3) is presented as Pearson’s R and Kendall’s *τ* . Bar charts in the right column (b, d) show the mean absolute error (MAE) between ground truth and predicted average CDR RMSDs, quantifying the magnitude of prediction errors for each method.

To contextualize these results, we introduce a CDRH3 Levenshtein distance baseline. Each antibody pair is assigned the mean RMSD of all test-set pairs sharing the same CDRH3 Levenshtein distance. The AbSLang models outperform this baseline, indicating that they capture structural information beyond what sequence similarity alone can provide.

### 3.2 AbSLang embeddings enable accurate structure similarity search

Antibody repertoire mining is the process of searching large libraries of antibodies for sequences with improved properties such as binding affinity, offering a route to antibody optimization without in vitro engineering [29]. Repertoire mining largely has been restricted to sequence similarity search or clonotyping. A structure-based similarity retrieval approach would significantly expand the coverage and effectiveness of repertoire mining.

We implemented AbSLang-search, a fast and memory efficient method for retrieval by structural similarity using embeddings generated by AbSLang. By indexing these structure-aware embeddings with FAISS [26], AbSLang-search rapidly retrieves antibodies within large databases, such as the Observed Antibody Space (OAS) [19], without requiring explicit 3D structure predictions and storage of PDB files.

We benchmarked AbSLang-search against an explicit structure-based retrieval approach using antibody structures predicted by ABodyBuilder3.

To evaluate whether AbSLang-search retrieves structurally similar antibodies, we used the AbSLang test set and computed CDR RMSD from experimental structures as the ground-truth ranking. For each query antibody, the true top-20 neighbors were defined as the 20 candidates with the lowest experimental-structure CDR RMSD to the query. We then compared the top-20 neighbors retrieved by AbSLang-search and by an ABodyBuilder3-predicted-structure baseline against this experimental ground truth.

Retrieval performance was assessed using: (1) top-20 overlap with the experimental-structure top-20 set, reported as the average number of correctly recovered neighbors out of 20; and (2) Kendall’s *τ* between the method-derived ranking and the experimental-structure ranking within the evaluated top-20 set. AbSLang-search achieves slightly lower but comparable overlap and rank correlation relative to ABodyBuilder3-based retrieval (Table 1).

**Table 1.** Retrieval performance against ground truth. For each query, the ground-truth top-20 set is defined using CDR RMSD computed from experimental structures. ‘Overlap’ is the average number of retrieved antibodies that are also in the ground-truth top-20 set. Kendall’s *τ* measures rank agreement with the ground truth ranking.

| **Paired** | AbSLang-Search | ABodyBuilder3 workflow |
| --- | --- | --- |
| Top 20 overlap | 13.60 | 14.40 |
| Top 20 $\tau$ | 0.25 | 0.30 |
| Average Precision Under 2 Å | 0.45 | 0.56 |
| **VH** | AbSLang-Search | ABodyBuilder3 workflow |
| Top 20 Overlap | 13.36 | 13.81 |
| Top 20 $\tau$ | 0.25 | 0.29 |
| Average Precision Under 2 Å | 0.55 | 0.58 |

We also evaluated distance-threshold retrieval, defined as the identification of antibody pairs with experimental CDR RMSD below 2 Å. In this setting, AbSLang-search again demonstrates comparable performance to the ABodyBuilder3-predicted-structure baseline, although with modestly lower average precision (Supplementary Figure 1).

### 3.3 AbSLang-Search is highly scalable across large databases

AbSLang-search significantly reduces storage and computational load as compared to explicit structure prediction. For example, storing ABodyBuilder3 predicted structures for databases such as paired-OAS (approximately 2.5 million sequences) would require around 508 GB of storage (Table 1). In contrast, our quantized embeddings can be stored within a FAISS index of just 879 MB. Moreover, searching the paired-OAS database with AbSLang-search takes merely 7 seconds, compared to approximately 18 minutes for the ABodyBuilder3-based approach.

We further benchmarked AbSLang-search specifically for heavy-chain sequences against KA-search, a rapid antibody sequence similarity search method [30], using a 10-million heavy chain sequence subset of the unpaired-OAS. AbSLang-search completes the query in 1.22 seconds, while KA-Search takes 8.3 seconds.

The computational efficiency of AbSLang-search which exceeds the speed of efficient sequence only methods makes the mining of large sequence databases by structural similarity tractable.

## 4 Conclusions

In this study, we present AbSLang, a model that generates sequence-level embeddings capturing antibody CDR structure similarity. AbSLang predicts pairwise CDR RMSDs with accuracy comparable to exact RMSD calculation from AlphaFold3 and ABodyBuilder3 predictions. We also introduce AbSLang-search which uses AbSLang embeddings and rapid vector database search tools to retrieve the top-K structurally similar antibodies from large sequence libraries. By enabling scalable structural search across resources such as OAS, AbSLang surfaces structurally similar antibodies with low sequence identity, accelerating therapeutic antibody discovery and repertoire-level studies. The AbSlang model and AbSLang-search pipeline are available on GitHub (https://github.com/ericjidawang/AbSLang/). We provide a web app which allows comparison of pairs of antibodies as well as searching OAS by structural similarity (www.opig.stats.ox.ac.uk/webapps/abslang).

## Notes

https://github.com/oxpig/AbSLang

### Competing Interest Statement

The authors have declared no competing interest.

