## Supplementary Information for "Fast retrieval of structurally similar antibodies from large sequence databases with AbSLang"

Eric Wang[1], Fabian C. Spoendlin[1], Alexander Greenshields-Watson[1], Christopher R. Taylor[1], and Charlotte M. Deane[1,*]

[1]Department of Statistics, University of Oxford, Oxford, UK

### 1 Supplementary figures

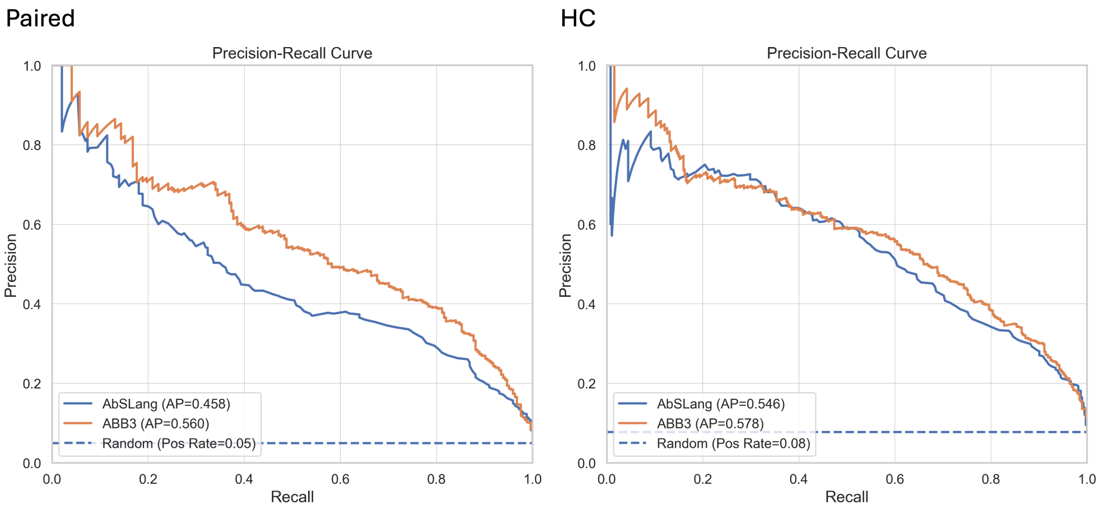


Figure 1: Precision-recall curves for retrieval of antibody pairs in heavy chain (HC) and paired sets with real RMSD < 2Å. AbSLang-search and the ABodyBuilder3-based workflow are compared to a baseline achieved by random retrieval of antibody pairs (Random).

### 2 VHH model

Despite similarities between camelid VHH IGHV genes and human antibody VH3 genes, VHH and antibody CDR loops adopt different conformations [1], due to longer CDR3s, absence of light chain (allowing the CDR3 to adopt a wider range of conformations) and non-canonical disulfide bonds between CDR loops[2]. Therefore, a separate model was trained apart from the antibody heavy chain specific model.

#### 2.1 Data and method

The overall model architecture and training method was carried over from the antibody specific versions (with 4 transformer encoder layers due to less VHH data being available to prevent overfitting). We replicate the training dataset from NanoBodyBuilder2 [3], resulting in 1,049 VHH structures. The same train/validation/test split was carried over from NanoBodyBuilder2.

### 2.2 Results

We benchmarked the VHH specific model against a state-of-the-art VHH specific structure prediction tool NanoBodyBuilder2, for retrieval and RMSD prediction accuracy like the antibody specific models.

For RMSD prediction, our model demonstrated comparable or superior performance relative to NanoBodyBuilder2 (Fig. 2). Additionally, our model achieved comparable error metrics to NanoBodyBuilder2, with both methods significantly outperforming a baseline model based on CDRH3 Levenshtein distance.

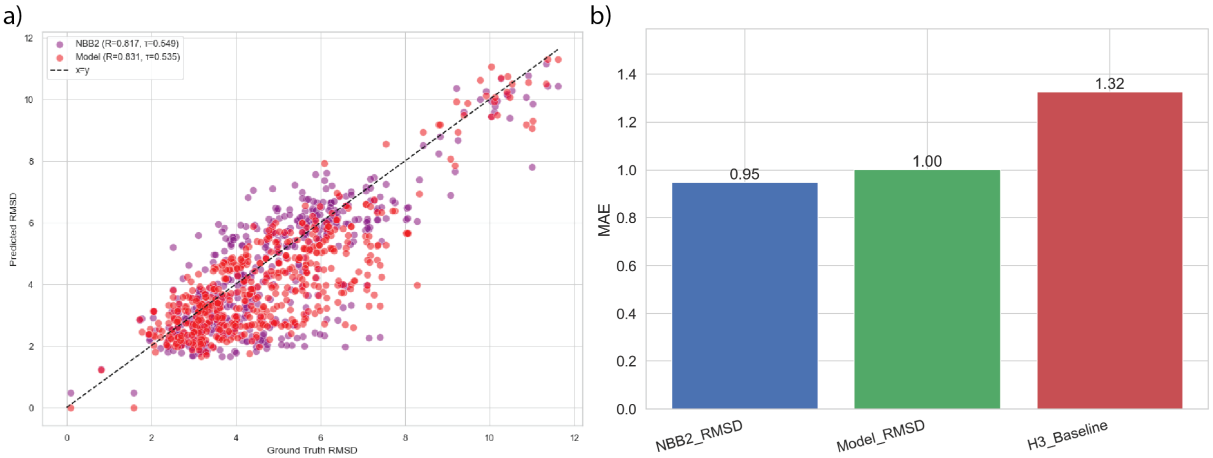


Figure 2: Performance of the VHH-specific model on raw CDR RMSD predictions. (a) Correlation between predicted and actual CDR RMSDs shows our model outperforming NanoBodyBuilder2 (NBB2) in overall correlation and exhibiting less prediction bias. NanoBodyBuilder2 tends to regress predictions towards known structures, resulting in underestimations of RMSDs. (b) Error comparison reveals our model performs similarly to NanoBodyBuilder and significantly outperforms a baseline predictor based on CDRH3 Levenshtein distances.

### 2.3 Retrieval Metrics of VHH Model

Instead of the 2.00 $\AA$ threshold used for HC and paired antibodies, a threshold of 2.5Å was chosen to assess retrieval accuracy for the VHH model due to differences in test set distributions. In the paired VH/VL and heavy-chain only antibody test sets, roughly 5% of antibody pairs fell below the 2Å RMSD threshold, while in the VHH test set, only 1.5% of pairs met this criterion. Thus, a higher threshold (2.5 Å) was selected to better reflect the structural similarity distribution in the VHH dataset.

We also assessed the precision and recall for the VHH model and NanoBodyBuilder2 for retrieval under 2.5 $\AA$.

Table 1: Retrieval performance against ground truth

| **Metric** | AbSlang-VHH-Search | NanoBodyBuilder2 workflow |
|---|---|---|
| Top 20 Overlap | 16.88 | 17.22 |
| Top 20 $\tau$ | 0.28 | 0.30 |
| Average Precision Under 2.5 $\AA$ | 0.32 | 0.29 |

[3] B. Abanades, W. K. Wong, F. Boyles, G. Georges, A. Bujotzek, and C. M. Deane, "ImmuneBuilder: Deep-Learning models for predicting the structures of immune proteins,"
